# Epididymal glucocorticoid receptors promote intergenerational transmission of paternal stress

**DOI:** 10.1101/321976

**Authors:** Jennifer C Chan, Bridget M Nugent, Kathleen E Morrison, Eldin Jašarević, Natarajan V Bhanu, Benjamin A Garcia, Tracy L Bale

**Affiliations:** Department of Biomedical Sciences, School of Veterinary Medicine and Perelman School of Medicine, University of Pennsylvania, Philadelphia, PA 19104, USA; Epigenetics Institute, Department of Biochemistry and Biophysics, Perelman School of Medicine, University of Pennsylvania, Philadelphia PA 19104, USA; Department of Pharmacology, Center for Epigenetic Research in Child Health and Brain Development, University of Maryland School of Medicine, Baltimore MD 21201, USA

## Abstract

Paternal preconception exposures and insults, including stress, dietary challenge and drugs of abuse, can shape offspring health and disease risk outcomes, as evidenced from retrospective human studies and more recent animal models^1–16^. Mechanistic examination has implicated small noncoding RNA populations in sperm, including microRNA (miRs), as carriers of paternal environmental information that consequently influence offspring development^15,17–21^. However, the cellular mechanisms by which these paternal signals are relayed to sperm and how they may persist remain unknown. Here, using our previously established paternal stress mouse model we identify caput epididymal epithelial glucocorticoid receptors as crucial upstream mediators of long-lasting germ cell programming. We show that glucocorticoid treatment of caput epididymal epithelial cells results in increased glucocorticoid receptor levels and enduring changes to the miR content of secreted extracellular vesicles (EVs), or epididymosomes, known to interact with sperm and alter their RNA content^22,23^. Further, significant changes were detected in the caput epididymal histone code long after stress ended, both *in vitro* and *in vivo*, as a potential mechanism whereby stress programmed enduring changes to EV miRs. Genetic targeting to reduce caput epididymal epithelial-specific glucocorticoid receptors reversed stress-induced chromatin remodeling and promoted cellular resilience to paternal stress, ultimately rescuing transmission of a stress dysregulated offspring phenotype. Taken together, these studies identify glucocorticoid receptor regulation of EV miRs in the caput epididymis as a key contributor in the intergenerational transmission of paternal environmental stress experiences.

## Main text

The contribution of preconception insults in the etiology of disease has garnered great interest in recent years, yet the crucial molecular mechanisms whereby an environmental insult is transmitted from somatic to germ cells and is able to persist long after the insult had ended, are not known. To address this, we utilized our established paternal stress mouse model in which we have previously demonstrated that stress-altered sperm miRs causally promote offspring brain reprogramming and stress dysregulation^6^, an endophenotype common to many neuropsychiatric disorders^24^. We focused on the contribution of glucocorticoids as an essential and necessary component of stress signaling that when elevated, bind and activate the low-affinity glucocorticoid receptor, a ubiquitously expressed molecule critical for the orchestration of cellular responses and chromatin remodeling^25,26^. Further, extracellular vesicles (EVs) from caput epididymal epithelial cells deliver important proteins, lipids, and RNAs, including miRs, to maturing sperm, altering sperm content^18,22,23,27,28^. Therefore, we hypothesized that in response to stress, caput epididymal epithelial glucocorticoid receptors are poised to contribute both to changes in the composition of the secreted EV miRs that interact with and shape maturing sperm, and also to coordinate local somatic epigenetic remodeling, as paternal-experienced stress produces effects that endure long after stress end^29^.

Therefore, to identify the molecular and epigenetic marks involved in the persistence of sperm miR alterations by stress, we administered chronic stress to male mice and collected epididymal sperm 1- or 12-weeks post-stress end (**Fig. 1a**, top) to compare the acute and enduring (allowing approximately two cycles of sperm turnover following stress exposure^30^) effects. We performed small RNA sequencing and identified two distinct populations of differentially expressed sperm miRs (adjusted *P* < 0.05), with no similarly stress-altered miRs shared between these populations (**Fig. 1b**), suggesting that a unique mechanism emerges following the acute response to stress to induce enduring changes in sperm miR populations.

**Figure 1.**
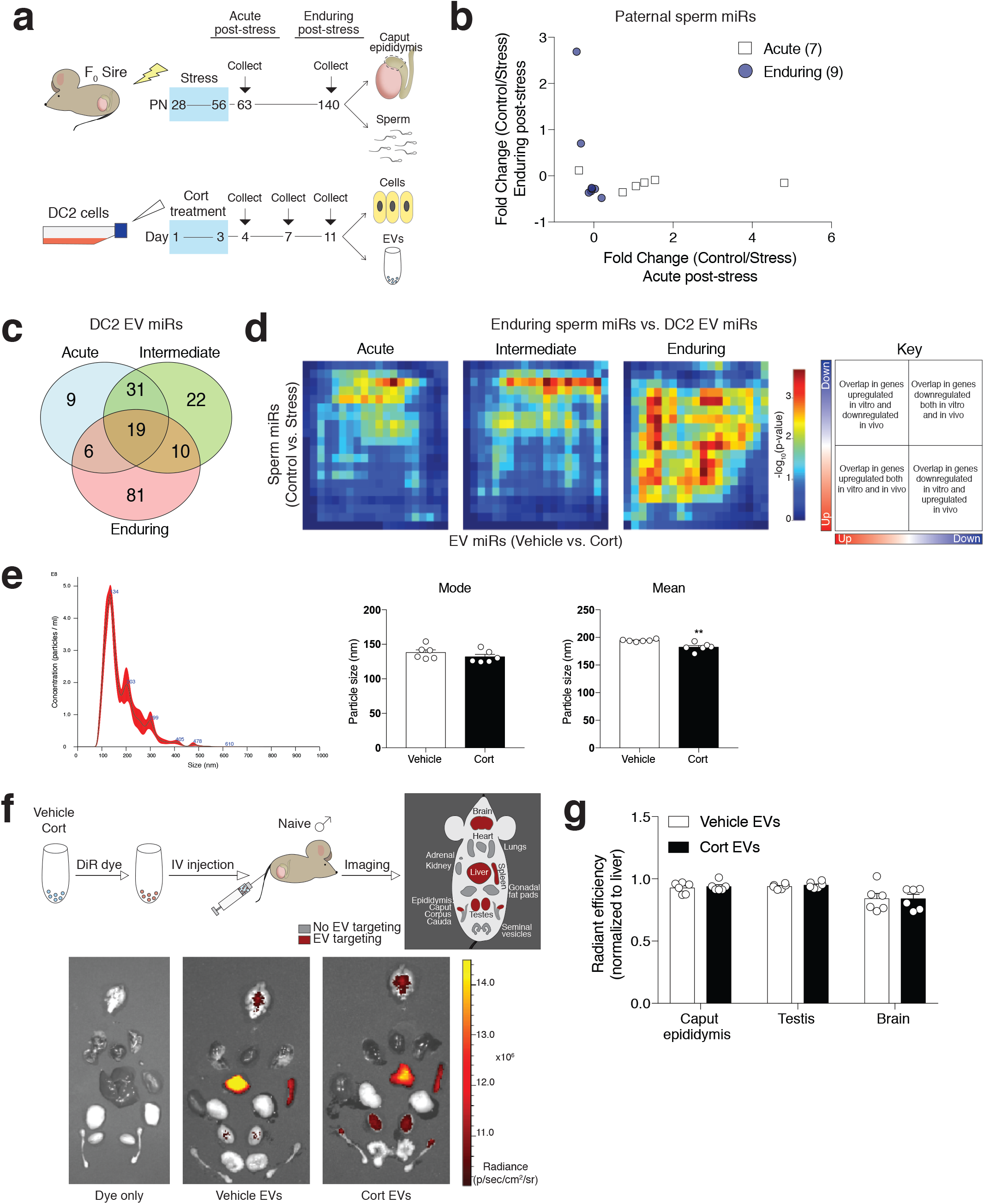
Glucocorticoid-treated DC2 mouse caput epididymal epithelial cell EVs *in vitro* mimic paternal stress programming of enduring sperm miRs *in vivo*. (**a**) Male mice were exposed to stress from postnatal days (PN) 28–56. Sperm and caput epididymal tissue were collected at 1-week (acute) and 12-weeks (enduring) post-stress. To mimic chronic stress, DC2 cells were administered three concentrations of corticosterone (cort) (50ng/ml (low), 500 ng/ml (medium), or 5 μg/ml (high)) for 3-days. Epididymal cells and secreted EVs isolated at 1 (acute), 4 (intermediate), and 8-days (enduring) post-treatment were examined for similar changes as those from paternal stress tissue. (**b**) Differential expression analysis of paternal sperm miRs identify distinct populations altered at 1- and 12-weeks post-stress, with each point representing one miR, suggesting unique mechanisms for acute and enduring sperm miRs post-stress. *N* = 6–8; adjusted *P* < 0.05. (**c**) Rank-rank hypergeometric overlap (RRHO) analysis was used between enduring *in vivo* paternal sperm miRs and *in vitro* DC2 EV miRs post-treatment to determine the best-matched period of miR regulation in DC2 cells. Venn diagram of significantly overlapping EV miRs from cells treated with the medium (physiologically relevant) corticosterone concentration following small RNA-sequencing, demonstrating the greatest overlapping number of miRs at 8-days post-treatment. *N* = 3–4; max –log_10_(*P*-value) = 3. These data are represented visually using (**d**) RRHO heatmaps where each pixel represents one miR comparison color-coded for degree of significance, with the most upregulated miRs at the bottom left corner and downregulated miRs at the top right corner (as described in the schematic, right). (**e**) Representative particle tracking plot (left) using Nanosight confirm DC2 EV size distribution. Corticosterone treatment did not affect the mode (Student’s t-test, t(10) = 1.165, *P* = 0.2712), but reduced the mean (Student’s t-test, t(10) = 3.865, *P* = 0.0031) of EV particle size 8-days post-treatment, suggesting altered lipid composition/function. *N* = 6. Student’s t-test, \*\**P* < 0.01. (**f**) Representative image of tissue-specific selectivity of 5E7 near-infrared DiR dye-labeled DC2 EVs treated with vehicle or corticosterone. (**g**) There were no differences between treatment for each tissue in total radiant efficiency of caput epididymis (Student’s t-test, t(10) = 0.4757, *P* = 0.6445), testes (Student’s t-test, t(10) = 0.8337, *P* = 0.4239), and brain (Student’s t-test, t(10) = 0.00912, *P* = 0.9929) between EVs injected, suggesting corticosterone-treated EVs retain endogenous tissue selectivity. *N* = 6. Data are mean ± SEM, with individual data points overlaid.

As we previously established that intergenerational transmission of paternal stress can continue for months after stress has ended, we investigated the epigenetic mechanism whereby epididymal EV miR changes, that are likely to impact sperm content during maturation, are maintained. To examine the specific population of caput epididymal EVs, we treated cultured DC2 mouse caput epididymal epithelial cells with corticosterone *in vitro*. Using DC2 cells allowed us to isolate EVs secreted into the media produced from a specific cell population. As all mammalian tissues secrete EVs into circulation, such select isolation *in vivo* is not possible^31^. Further, this allowed for the controlled administration of corticosterone, the primary glucocorticoid in rodents produced by activation of the hypothalamus-pituitary-adrenal (HPA) stress axis and known to activate the low affinity glucocorticoid receptors^25^. We confirmed the purity of EVs isolated from DC2 cells using validated EV markers^32^ (**Supplementary Fig. 1**). To develop an accurate modeling of the timing of events of this ‘stress in a dish’ model compared to our *in vivo* paternal model, we first examined three distinct concentrations of corticosterone that included the range of the mouse physiological baseline (low) and stress response (medium), as well as a supraphysiological (high) concentrations, at three time points post-treatment to examine the acute, intermediate, and enduring effects of treatment in DC2 cells (**Fig. 1a**, bottom). We then used rank-rank hypergeometric overlap (RRHO) analyses to evaluate the extent of overlap between stress- and corticosterone-altered miRs *in vivo* and *in vitro*, respectively. We compared control v. stress enduring differential expression profiles in sperm miRs and the vehicle v. corticosterone differential expression profiles in DC2 EV miRs at each collection point post-treatment allowing for threshold-free identification followed by quantification of statistically significant overlap between datasets^33^. Using this approach, we compared the populations of significantly overlapping EV miRs following small RNA sequencing and found distinct groups of altered EV miRs that were dependent on the time post-treatment, where the degree of overlap increased dramatically at 8-days compared to 1-day post-treatment at all corticosterone concentrations (**Fig. 1c** and **Supplementary Fig. 2a**), supporting enduring effects present in our *in vitro* model. Following quantification of total overlapping EV miRs at all corticosterone concentrations and times, we confirmed that 8-days following treatment with the stress-relevant concentration of corticosterone most-closely matched the *in vivo* enduring paternal stress sperm (**Fig. 1d** and **Supplementary Fig. 2b**), where the total proportion of significantly overlapping miRs rose to 31.4 % (116/369). No doubt, the complexity of sperm miR composition reflects additional interactions from along the reproductive tract and therefore will not completely mirror the DC2 EVs, as has been described^28,34^.

To ensure corticosterone treatment did not disrupt the endogenous properties and tissue selectivity of DC2 EVs *in vivo*, we quantified and characterized the size distribution of DC2 EVs using Nanosight particle tracking analysis. Interestingly, corticosterone treatment reduced EV mean size, but not mode, consistent with possible changes to lipid or protein composition that may affect EV performance at select tissues^35^ (**Fig. 1e**). We labeled and isolated vehicle- and corticosterone-treated DC2 EVs with a near-infrared, lipophilic DiR dye, and injected 50 million EVs intravenously into naïve male mice (**Fig. 1f**, top schematic). 24-hours post-injection, we imaged the tissues to evaluate the bio-distribution of caput epididymal EV targeting. As expected, there was substantial accumulation of EVs in the liver and spleen, as previously described for EVs from most other cellular sources^36^. However, specific to EVs from these epididymal epithelial DC2 cells, there was substantial accumulation along the reproductive tract, including the caput epididymis and testes, and a surprising localization to the brain (**Fig. 1f**). Importantly, corticosterone treatment of DC2 cells did not alter EV tissue targeting (**Fig. 1g** and **Supplementary Fig. 3**). These results suggest that stress at the level of the caput epididymis impacts EV and sperm miR content without disruption to endogenous EV tissue selectivity. The local effects of EV miRs on paternal tissues such as the brain remain to be evaluated.

To assess the role of glucocorticoid receptors in the prolonged timing effects of EV miRs, we performed immunoblotting on DC2 cells 1- and 8-days following corticosterone treatment. While there were no significant changes in glucocorticoid receptor levels immediately following corticosterone treatment end (**Fig. 2a**, left), glucocorticoid receptor levels were increased 8-days post-treatment at all corticosterone concentrations (**Fig. 2a**, right). We hypothesized that increased nuclear glucocorticoid receptors may coordinate chromatin remodeling to promote enduring changes to EV miR content. As changes to histone composition and post-translational modifications (PTM) are a likely candidate for upstream broad transcriptional control of miR genes, we performed unbiased quantitative histone mass spectrometry in DC2 cells 8-days post-corticosterone treatment (**Supplementary Table 1**). We applied Random Forests classification analysis to our dataset to identify the histones and PTMs altered by corticosterone. Random Forests is an ensemble-learning algorithm that identifies groups of features (i.e. histone PTMs) altered together, and ranks these features according to their importance to the model’s accuracy^37^. Using this approach, we identified the top thirteen histone PTMs, as determined by tenfold cross-validation of the model (**Fig. 2b**, inset), that most accurately discriminate vehicle v. corticosterone-treated DC2 cells (**Fig. 2b**). To confirm these Random Forests results, we performed Mann-Whitney U tests (**Fig. 2c**), demonstrating long-term remodeling of the histone code that corresponded with post-treatment glucocorticoid receptor increases.

**Figure 2.**
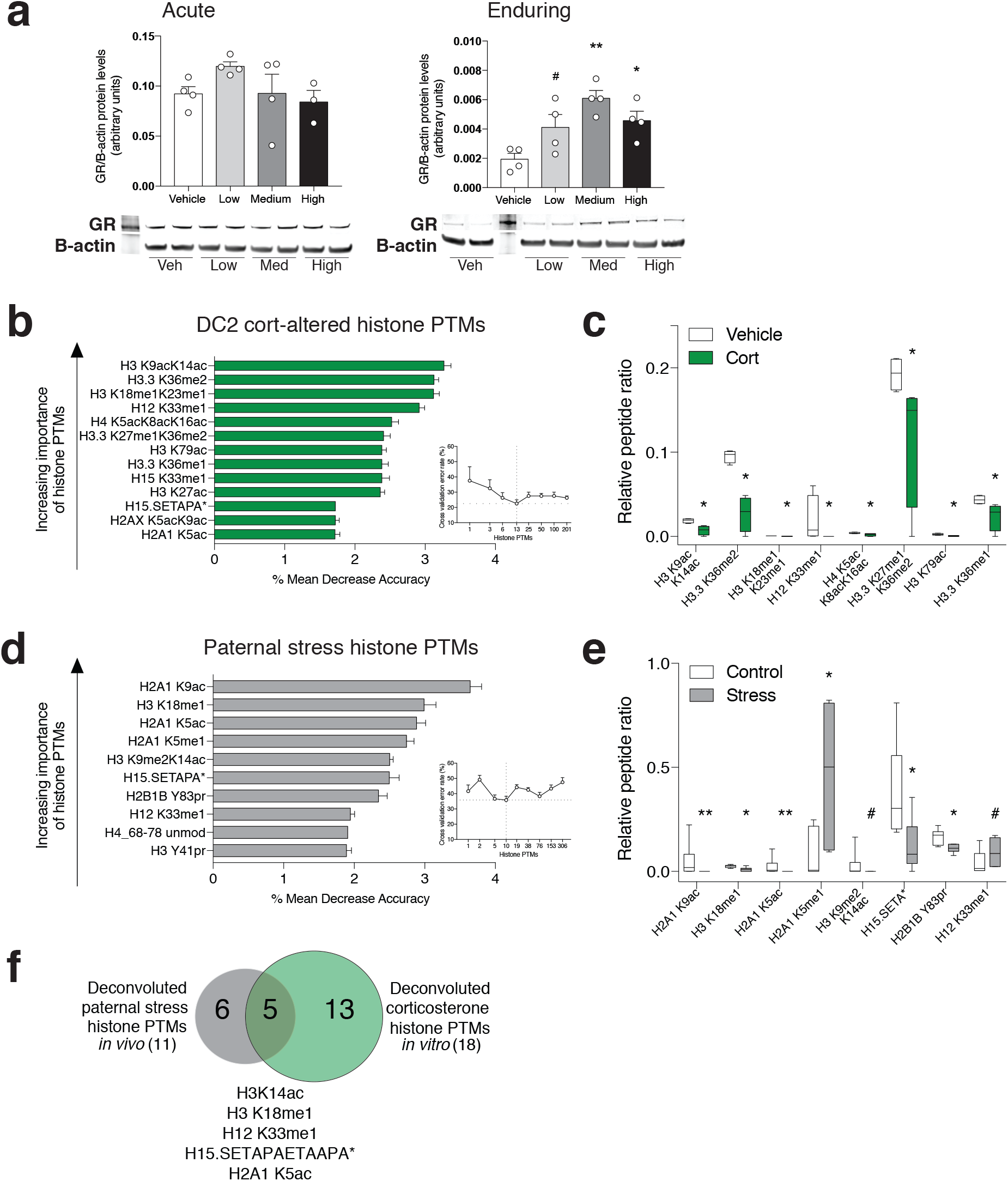
Glucocorticoid receptors are increased post-stress and correspond with enduring reprogramming of the caput epididymal histone code. (**a**) Immunoblotting of glucocorticoid receptor levels 1- (acute time point, left) and 8-days (enduring time point, right) post-treatment. No effect of corticosterone treatment at the acute time point (one-way ANOVA, F(3, 11) = 1.644, *P* = 0.2360). There were significant treatment effects at the enduring time point (one-way ANOVA, F(3,12) = 7.306, *P* = 0.0048. Bonferroni’s post-hoc analysis showed significant differences between vehicle (Veh) v. medium (Med) concentration (t(12) = 4.625, adjusted *P* = 0.0018); vehicle v. high concentration (t(12) = 2.93, adjusted *P* = 0.0378), and a nonsignificant difference between vehicle v. low concentration (t(12) = 2.416, adjusted *P* = 0.0977), suggesting glucocorticoid receptors (GR) are involved in enduring EV miR alterations. *N* = 3–4; one-way ANOVA with Bonferroni’s correction, \*\**P* < 0.01, \**P* < 0.05, #*P* < 0.1. Data are mean ± SEM, with individual data points overlaid. (**b** and **d**) Random Forests analysis of quantitative histone post-translational modifications (PTM) mass spectrometry identified the top histone PTMs, ranked by importance, that most accurately discriminate at the enduring time point between (**b**) vehicle v. corticosterone treatment of DC2 cells *in vitro*, and (**d**) control v. stress caput epididymis *in vivo*. Mean decrease accuracy indicates the percent decrease in model accuracy if the histone PTM is removed. *N* = 4–6. Error bars are ± SD. (**c** and **e**) Relative abundance of the top eight histone PTMs determined by Random Forests were confirmed by Mann-Whitney U tests between treatment for each individual histone PTM in (**c**) DC2 cells and (**e**) paternal stress caput epididymis. Data are median ± interquartile range. *\*\*P* < 0.01, \**P* < 0.05, #*P* < 0.1. (**f**) Venn diagram of total deconvoluted histone PTMs discriminating treatment groups, as determined by Random Forests analysis, between *in vivo* caput epididymis (gray) and *in vitro* DC2 cells (green), and their overlap (histone PTMs listed below).

We then compared these enduring *in vitro* DC2 epigenetic changes to our *in vivo* paternal stress model. We again performed histone PTM mass spectrometry on whole caput epididymal tissue from control and stress males 12-weeks post-stress end, and used Random Forests analyses (**Supplementary Table 2**). We identified ten histone PTMs that most accurately classified our model (**Fig. 2d**), and that were substantiated by Mann-Whitney U tests (**Fig. 2e**). We deconvoluted these histone PTMs identified by Random Forests in our *in vitro* and *in vivo* models and identified five overlapping histone PTMs (**Fig. 2f**), approximately 45% (5/11) of total treatment-discriminating *in vivo* histone PTMs, supporting that our paternal ‘stress in a dish’ model, where glucocorticoids were administered, extensively mimics features of endogenous paternal stress programming. These five common histone PTMs include modifications to two H1 variants (H12 and H15), H2A1 K5 acetylation, H3 K18 monomethylation, and H3 K14 acetylation. Interestingly, H3 K14ac has been implicated in driving stress effects at the chromatin level in other stress models^38–40^. While the literature is scarce regarding the remaining histone PTMs, these data suggest that post-stress glucocorticoid receptor increases may mediate chromatin remodeling at specific loci, including those of EV miRs, to alter their expression, consistent with previous reports^41–43^. Moreover, these data support that stress in the environment is able to promote lasting modifications to cellular transcriptional machinery within reproductive tissues, functionally modifying germ cell content.

To then examine a causal role of epididymal epithelial glucocorticoid receptors in the intergenerational transmission of paternal stress *in vivo*, we genetically targeted glucocorticoid receptors in male mice to reduce expression (GR^Het^) specifically in caput epididymal epithelial cells using the lipocalin-5 (Lcn5) promoter^44^ crossed with GR^flox^ mice^45^ (**Fig. 3a**). We additionally incorporated the transgenic RiboTag line, allowing isolation of mRNA specifically from the HA-tagged ribosomal subunit, Rpl22, in caput epididymal epithelial cells for RNA sequencing^46^. In these mice, we verified transgenic reduction as well as the inhibition of stress-mediated increases in glucocorticoid receptors 12-weeks post-stress in GR^Het^ mice (**Fig. 3b**). We hypothesized that inhibiting post-stress glucocorticoid receptor increases here would prevent the enduring intergenerational transmission of the paternal stress phenotype. To test this, we bred control and stressed GR^WT^ and GR^Het^ males to wildtype females, and examined the offspring HPA stress axis response. Remarkably, there was a significant paternal treatment x paternal genotype interaction in the offspring response to an acute restraint whereby paternal epididymal GR^Het^ prevented the blunted offspring stress response to control levels (**Fig. 3c**). We extended this finding by examining the effect in response to an additional type of HPA activation, an acute predator odor, as a more ethologically relevant challenge in mice. Again, paternal stress GR^WT^ offspring presented with a dysregulated HPA axis response compared to paternal control offspring, and this heightened response was again rescued in paternal stress GR^Het^ offspring (**Fig. 3d**). Importantly, neither treatment nor genotype affected paternal reproductive function or litter characteristics (**Supplementary Table 3**). The paraventricular nucleus (PVN) of the hypothalamus is key to HPA stress regulation and we have previously demonstrated transcriptional dysregulation of the PVN in paternal stress offspring^6^. Therefore, we next examined the ability of the GR^Het^ paternal genotype to rescue these gene expression changes in the offspring PVN. As expected, hierarchical clustering of all genes altered by paternal stress exposure demonstrated that the greatest difference in PVN gene expression was between control and paternal stress GR^WT^ offspring, supporting a strong programming effect of paternal stress, while paternal GR^Het^ mitigated the extent of PVN changes by paternal stress (**Fig. 3e**), demonstrating that caput epididymal glucocorticoid receptors govern paternal stress transmission of the offspring brain and stress response.

**Figure 3.**
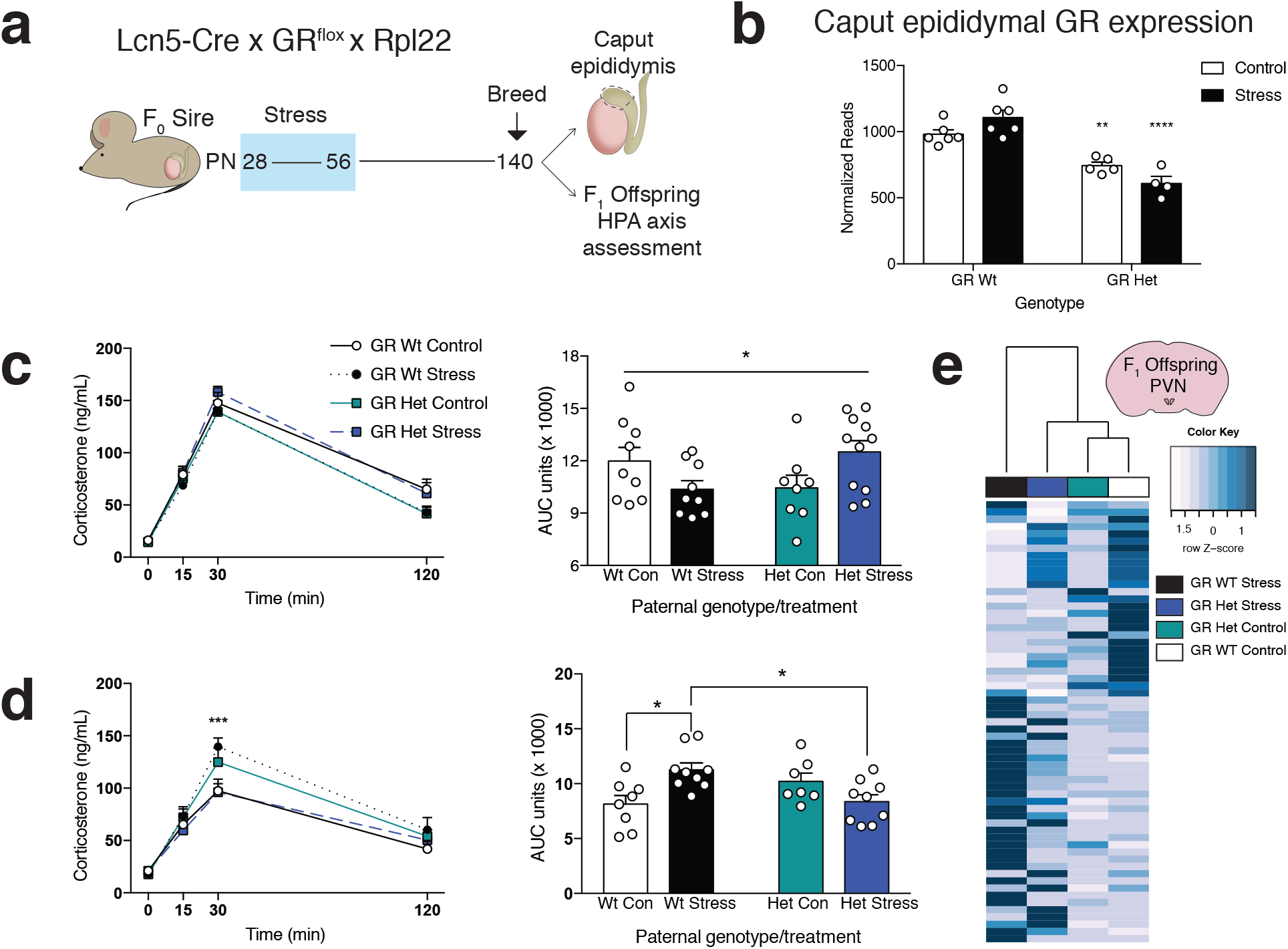
Genetic reduction of caput epididymal epithelial glucocorticoid receptors *in vivo* rescues paternal stress programming of offspring stress dysregulation. (**a**) Caput epididymal epithelial cell-specific Lcn5-Cre x GR^flox^ x Ribotag (Rpl22) male mice were exposed to stress, as above, and were bred 12-weeks post-stress. Adult offspring were assessed for HPA stress axis responsivity. (**b**) To ensure transgenic glucocorticoid receptor (GR) reduction and inhibition of post-stress glucocorticoid receptor increases, glucocorticoid receptor mRNA expression from paternal caput epididymal epithelial cells was examined using Ribotag technology (two-way ANOVA, main effect of genotype (F(1, 17) = 68.71, *P* < 0.0001), interaction of genotype x treatment (F(1,17) = 8.652, *P* = 0.0091). Tukey’s post-hoc analysis showed significant differences between Control GR^WT^ and Control GR^Het^ (t(17) = 5.527, adjusted *P* = 0.0056) and between Stress GR^WT^ and Stress GR^Het^ (t(17) = 10.89, adjusted *P* < 0.0001)). *N* = 4–6; Tukey’s post-hoc test, \*\**P* < 0.01, \*\*\*\**P* < 0.0001. (**c**) There was a significant interaction for paternal treatment x genotype for the offspring HPA area under the curve (AUC), where the reduced response to an acute restraint in wildtype (Wt) paternal stress offspring was normalized by paternal glucocorticoid receptor reduction (two-way ANOVA, interaction of paternal genotype x paternal treatment, F(1, 34) = 4.902, *P* = 0.0336). *N* = 8–11; two-way ANOVA, \**P* < 0.05. (**d**) Similarly, there was a significant interaction in the AUC (two-way ANOVA, interaction of paternal genotype x paternal treatment, F(1, 29) = 12.65, *P* = 0.0013. Tukey’s post-hoc analysis showed significant differences between Control GR^WT^ offspring v. Stress GR^WT^ offspring (t(29) = 4.554, adjusted *P* = 0.0158) and between Stress GR^WT^ offspring v. Stress GR^Het^ offspring (t(29) = 4.369, adjusted *P* = 0.0216)) and a main effect on the curve (two-way ANOVA with time as a repeated measure, main effect of treatment (F(3, 29) = 3.325, *P* = 0.0333) and main effect of time (F(3, 87) = 97.71, *P* < 0.0001). Tukey’s post-hoc analysis showed significant differences at the 30-minute time point between Control GR^WT^ offspring v. Stress GR^WT^ offspring (t(116) = 5.183, adjusted *P* = 0.0021) and between Stress GR^WT^ offspring v. Stress GR^Het^ offspring (t(116) = 5.125, adjusted *P* = 0.0009)), whereby paternal GR^Het^ prevented the paternal stress-altered HPA response to an acute predator odor exposure in offspring. *N* = 7–9; Tukey’s post-hoc test on the curve, \*\*\**P* < 0.001; Tukey’s post-hoc test on the AUC, *\*P* < 0.05. Data are mean ± SEM, with individual data points overlaid. (**e**) Hierarchical clustering and heatmap of all differentially expressed genes between paternal stress and control groups from RNA-sequencing of the paraventricular nucleus (PVN) from naïve adult offspring, showing caput epididymal GR^Het^ mitigation of paternal stress programming. *N* = 5–6; adjusted *P* < 0.05.

To then determine the mechanism within the caput epididymis that can promote or prevent enduring transmission, we performed differential expression analyses on the actively translating genes isolated from HA-tagged Rpl22 subunits (RiboTag) in Lcn5 + cells. Remarkably, comparing within genotype for the effects of stress, there were 1826 differentially expressed genes (adjusted *P* < 0.05) affected by stress 12-weeks following stress-end between GR^Het^ mice, but very few (65 genes) between GR^WT^ mice (**Fig. 4a**), suggesting that reducing caput glucocorticoid receptors results in a robust, compensatory response to stress within these epididymal cells. Importantly, there were 176 differentially expressed genes between control mice and 810 genes between stress mice; therefore, the robust response of GR^Het^ mice to stress was not attributed to glucocorticoid receptor reduction alone. In comparison, using the same pipeline, there was a modest caput epididymal response acutely post-stress, where the total number of genes altered (adjusted *P* < 0.05) in all comparisons totaled 62 (**Fig. 4b**). Comparing the number of acute v. enduring stress-altered differentially expressed genes, there was a 3-fold induction in GR^WT^ males and a 200-fold induction in GR^Het^ males (**Fig. 4c**), supporting the time post-stress as a crucial window whereby stress is processed to promote long-term transmission. Lastly, to determine the functional pathways broadly affected by the interaction of treatment and genotype in the caput epididymis that remain altered long-term, we performed cluster analyses and identified three clearly distinct groups of co-regulated genes (**Fig. 4a**, heatmap side). For each cluster, we used ClueGO for functional annotation analysis and gene ontology (GO) terms for biological processes to inform us as to pathways that may be driven by stress, by genotype, or by both^47^. Genes from cluster 1 were clearly changed specifically in control GR^Het^ mice, an effect driven by a reduction in glucocorticoid receptors alone, and enriched for GO terms including cell-cell signaling and vesicle-mediated transport (**Supplementary Table 4**), suggesting caput glucocorticoid receptors normally regulate epithelial cell communication with other cell types. Related to our hypothesis, cluster 2 genes were upregulated in GR^WT^ mice by prior stress experience, and intriguingly, were reversed in GR^Het^ stressed mice, supporting that this cluster includes genes involved in enduring programming of intergenerational transmission. Genes from cluster 2 were most significantly enriched for GO terms representing chromatin-modifying processes and intracellular transport (**Fig. 4d**, left, and **Supplementary Table 5**), again corroborating that stress reprograms the histone code long-term. Cluster 3 genes were upregulated specifically in GR^Het^ mice exposed to prior stress, and were enriched for ribosomal biogenesis, mitochondrial transport and metabolic processes (**Fig. 4d**, right, and **Supplementary Table 6**), suggesting increased oxidative phosphorylation capacity by mito-ribosome biogenesis may be a counteractive response to stress^48^. Altogether, these data indicate that paternal stress transmission is glucocorticoid receptor-dependent, where caput epididymal glucocorticoid receptor reduction reverses stress-induced chromatin remodeling and enhances mitochondrial function, promoting cellular resilience to environmental challenges^49^ and preventing transmission of an offspring phenotype.

**Figure 4.**
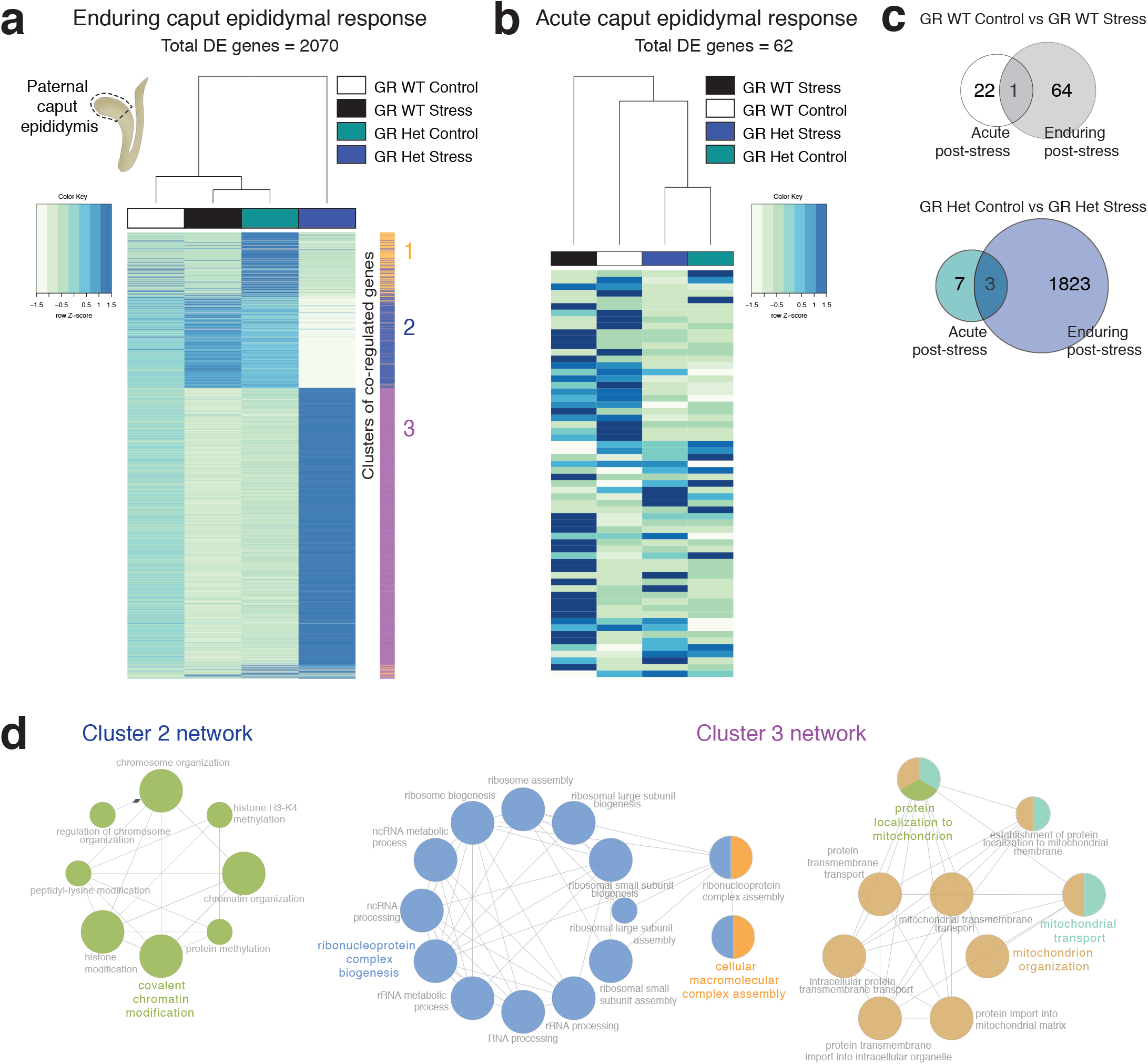
Reduction of caput epididymal glucocorticoid receptors reverses stress-induced epigenetic programming and promotes ribosomal and mitochondrial processes. (**a, b**) Heatmap of all differentially expressed (DE) genes from RNA-sequencing of paternal caput epididymal epithelial cells isolated using Ribotag technology at (**a**) 12- and (**b**) 1-week post-stress. *N* = 4–6; adjusted *P* < 0.05. Hierarchical clustering of co-regulated genes for (**a**) is depicted by color blocking on right of heatmap for functional annotation analysis. (**c**) Venn diagrams of the acute v. enduring caput epididymal epithelial response to prior stress exposure between GR^WT^ males (**top**) and GR^Het^ males (**bottom**), substantiating a post-stress mechanism that mediates enduring changes. (**d**) Functional annotation analysis of the enduring caput epididymal response 12-weeks post-stress using gene ontology terms (biological processes) for cluster 2 (genes increased in Stress GR^WT^ and decreased in Stress GR^Het^, **left**) and cluster 3 (genes increased only in Stress GR^Het^, **right**) reveal pertinent enriched pathways, determined by ClueGo and depicted as a network. Edges indicate degree of connectivity between terms. Node size indicates statistical significance, with the leading term (large, colored descriptor) determined by greatest degree of significance. Node colors indicate number of groups associated with the gene ontology term.

In summary, these studies identify a cellular mechanism whereby paternal stress experience produces lasting consequences for future offspring neurodevelopment. Our findings implicate caput epididymal epithelial glucocorticoid receptors as important orchestrator of environmental stress contributing to enduring epigenetic reprogramming and epididymal epithelial EV and sperm miR changes. We show that caput epididymal glucocorticoid receptor reduction coordinates a compensatory response to stress, including reversal of chromatin modifications that ultimately rescues paternal stress transmission of the offspring phenotype. These studies establish the paternal caput epididymis as a key determinant in the intergenerational transmission of environmental experience and programming of offspring disease risk.

## Materials and Methods

**Animals**. Male C57BL/6J and female 129S1/SvImJ mice were obtained from Jackson Laboratories and were used to produce C57BL/6:129 F1 hybrids. F1 hybrids were used for all paternal stress studies. For the caput epididymal epithelial cell-specific reduction of GR and RiboTag breedings, GR^flox^ (B6.129S6-*Nr3c1^tm2.1Ljm^/*J) and RiboTag (B6N.129-*Rpl22^tm1.1Psam^*/J) mice were crossed with 129S1/SvImJ females for minimally 3 generations^50^. Lcn5-Cre male mice on a C57Bl/6J background were purchased from the Model Animal Research Center of Nanjing University and were bred to double heterozygous GR^flox^; RiboTag 129 females to generate experimental animals. All mice were housed in a 12:12 light:dark cycle with temperature 22°C and relative humidity 42%. Food (Purina Rodent Chow; 28.1% protein, 59.8% carbohydrate, 12.1% fat) and water were provided ad libitum. All studies were performed according to experimental protocols approved by the University of Pennsylvania Institutional Animal Care and Use Committee, and all procedures were conducted in accordance with the NIH Guide for the Care and Use of Laboratory Animals.

**Chronic Variable Stress**. Administration of chronic variable stress was performed as previously described^6^. At PN28, males were weaned, pair-housed with a same-sex littermate, and randomly assigned to a control or stress group. Chronic variable stress occurred over 28 days (PN28–56). One stressor was administered each day and the order of stressors was randomized each week. Stressors include the following: 36 h constant light, 1 h exposure to predator odor (1:5000 2,4,5-trimethylthiazole (Acros Organics) or 1:2000 phenethylamine (Sigma)), 15 min restraint, novel object (marbles or glass vials) overnight, multiple cage changes, 100 dB white noise overnight, and saturated bedding overnight.

**Breeding**. Following completion of chronic variable stress (PN56), males were all left undisturbed for at least 1 week to remove the acute effects of stress. Males were then housed with virgin, stress-naïve F1 hybrid females at either 1- or 12-weeks following the end of stress exposure for a maximum of 3 nights. To minimize male-female interactions that may impact maternal investment or care^51^, observation of a copulation plug within 1 h after lights on signaled the immediate removal of the female to her own cage containing a nestlet.

**Adult tissue collection**. Sires were rapidly decapitated under isoflurane anesthesia 24 h following copulation. The testes, caput and corpus epididymis were removed and flash frozen in liquid nitrogen. Sperm were obtained by mincing the caudal epididymis into 1% BSA and subsequently isolated at 37°C through a double swim-up assay. The supernatant containing motile sperm was centrifuged for 5 min at 4000 rpm and the sperm pellets were stored at –80°C. Adult offspring were dissected at ∼20 weeks of age. Whole brains were removed, frozen on dry ice, and stored at –80°C.

**HPA axis assessment**. Plasma corticosterone was measured in response to an acute 15 min restraint stress in a 50mL conical tube. Testing occurred 2–5 h after lights on. Tail blood was collected at onset and completion of restraint (0 and 15 min) and 15 and 115 min after the end of restraint (30 and 120 min). Samples were immediately mixed with 50 mM EDTA and centrifuged 10 min at 5000 rpm. 3ul of plasma was collected at stored at –80°C until analysis. Corticosterone levels were determined by ^125^I-corticosterone radioimmunoassay (MP Biomedical) according to manufacturer’s protocol. For HPA axis responsivity to fox odor exposure, 1:5000 2,4,5-trimethylthiazole (Acros Organics) was administered on a Q-tip cotton swab in a separate testing room for 15 min to minimize odor exposure during recovery. For each experiment, no more than two littermates were included in each group.

**Cell culture and corticosterone treatment**. Immortalized mouse distal caput epididymal epithelial (DC2) cells were purchased from Applied Biological Materials and cultured as previously described^52^. Briefly, cells were seeded in 75cm^2^ Nunc EasYFlasks (Thermo Fisher) coated in collagen type 1, rat tail (Millipore). Cells were grown in Iscove’s Modified Dulbecco’s Medium (IMDM) supplemented with 10% exosome-free fetal bovine serum (Gibco) and 1% penicillin-streptomycin (Gibco). At monolayer confluency, the media was replaced and cells were either treated with 1:1000 vehicle (ethanol; resulting in 0.1% ethanol) or 1:1000 corticosterone in ethanol (Sigma; low concentration 144μM, medium concentration 1.4mM, high concentration 14.4mM –resulting in about 50, 500, or 5000 ng/ml of corticosterone, respectively). Cells were treated every 24 h for 3 days for a total of three treatments. The media was replaced 24 and 96 h following the last treatment. Media and cells were collected at 24, 96, or 192 h following the last treatment. For cell collection, cells were trypsinized in 0.25% trypsin-EDTA (Gibco), centrifuged at 1500 rpm for 3 min, and frozen at –80°C until further analysis.

**Extracellular vesicle (EV) isolation**. EVs were isolated from exosome-free media (Gibco) using differential centrifugation^32^. Briefly, cellular debris was removed from the media by centrifugation at 200g for 10 min, 2000g for 10 min, and 10,000g for 30 min. EVs were pelleted by ultracentrifugation at 100,000g for 1 h using the Optima L-90K Ultracentrifuge and SW 32 Ti swinging bucket rotor (Beckman Coulter). The EV pellet was resuspended in PBS or TriZol reagent and frozen at –80°C until further analysis.

**Nanoparticle tracking analysis**. All samples were run on a NanoSight NS500 to determine the size distribution of EV particles at the Center for Nanotechnology in Drug Delivery at the University of North Carolina. All samples were diluted to a concentration between 1EE08–5EE08 particles/mL in filtered PBS. Five 40 sec videos were taken of each sample to capture particles moving by way of Brownian motion. The nanosight software tracked the particles individually and using the Stokes-Einstein equation, calculated the hydrodynamic diameters.

**IVIS Spectrum Imaging of labeled EVs**. EVs isolated 8 days following 3 day treatment were labeled with XenoLight DiR Fluorescent Dye (PerkinElmer) per manufacturer’s instruction. Briefly, EV pellets were resuspended in 600 μl cold PBS and incubated with 20 μl 10mM DiR dye for 5 min at RT. As a non-EV control, 600 μl PBS alone was processed in parallel. The total volume was brought up to 38 ml with PBS and ultracentrifuged at 100,000g for 1 h. The dyed EV pellet was resuspended in PBS and 5e7 particles were injected intravenously via the tail vein into naïve adult F1 hybrid male mice. 24 h following injection, the mice were sacrificed and their tissues were collected for imaging using an IVIS Spectrum (PerkinElmer). The excitation filter was set at 745 and the emission filter was set at 800. For quantification, total radiant efficiency was calculated using Living Image software, with the minimum set at 1e7 and the maximum set at 1.45e7. Total radiant efficiencies for each tissue were normalized to total radiant efficiency of 0.1 g liver to control for success of the injection.

**Protein extraction and western immunoblotting**. Cell pellets were processed for immunoblotting using established protocols. For nuclear extractions, samples were homogenized with a pestle in cold sterile PBS, homogenates were centrifuged at 1200 g for 10 min at 4°C, pellets were washed with PBS, and resuspended in Buffer A (10mM Hepes pH 7.8, 10 mM MgCl_2_, 0.1 mM EDTA, 1 mM DTT, protease inhibitor cocktail (Sigma), phosphatase inhibitor cocktail (Sigma)). Following a 15 min incubation on ice, 0.05% NP-40 was added, samples were vortexed, and nuclear extracts pelleted at 14,000 x g for 30 sec. Nuclear pellets were resuspended in Buffer B (50 mM Hepes pH 7.8, 50 mM KCl, 300 mM NaCl, 0.1 mM EDTA, 1 mM DTT, protease inhibitor cocktail, phosphatase inhibitor cocktail). For whole cell extracts and EV protein extraction, samples were homogenized and resuspended in radioimmunoprecipitation assay (RIPA) buffer with protease inhibitor cocktail (Sigma), rotated for 2 h at 4°C, and pelleted at 5000 x g for 10 min. Protein quantification was done using Bradford assay (BioRad). For immunoblotting, twenty μg of protein was loaded per lane for gel electrophoresis onto a NuPAGE 4–12% Bis-Tris gel (Life Technologies). After running, gels were cut and the same molecular weight sections for all samples were transferred together to enable multiple probing and to control for transfer conditions. After transfer of proteins to a nitrocellulose membrane (Life Technologies), membranes were blocked with Odyssey blocking buffer (Li-Cor) and probed with rabbit anti-GR (1:10000; Abcam ab109022), mouse anti-beta actin (1:30000; Sigma A5441), rabbit anti-CD63 (1:1000; Systems Biosciences EXOAB-CD63A-1), rabbit anti-Lamp1 (1:1000; Abcam ab22595), and/or rabbit anti-Calnexin (1:1000; Abcam ab24170), followed by incubation in IRDye800-conjugated donkey anti-rabbit secondary (1:20,000; Li-Cor) and/or IRDye680-conjugated goat anti-mouse secondary (1:20,000; Li-Cor).

**Histone extraction, bottom-up nanoLC MS/MS and data analysis**. Samples were processed as previously described^53^. Briefly, whole caput epididymides or DC2 cell pellets were homogenized in nuclei isolation buffer (15mM Tris-HCl pH 7.5, 60 mM KCl, 15mM NaCl, 5mM MgCl2, 1 mM CaCl2, 250 mM sucrose) with 1 mM DTT, 1% phosphatase inhibitor (Sigma), 1 pellet protease inhibitor (Roche), 10mM sodium butyrate (Sigma), and 10% NP-40. Histones were acid extracted from nuclei by rotating overnight in 0.4N H2SO4 at 4°C and precipitated with 100% trichloroacetic acid overnight at 4°C. Extracted histones were washed with acetone and quantified by Bradford reagent according to manufacturer’s protocol (Sigma). ∼20ug histones were derivatized using propionic anhydride (Sigma) and digested with 1:10 trypsin (Promega). Samples were subsequently desalted by binding to C18 material from a solid phase extraction disk (Empore), washed with 0.5% acetic acid, and eluted in 75% acetonitrile and 5% acetic acid. Peptides were separated in EASY-nLC nanoHPLC (Thermo Scientific, Odense, Denmark) through a 75 μm ID x 17 cm Reprosil-Pur C_18_-AQ column (3 μm; Dr. Maisch GmbH, Germany) using a gradient of 0–35% solvent B (A = 0.1% formic acid; B = 95% acetonitrile, 0.1% formic acid) over 40 min and from 34% to 100% solvent B in 7 minutes at a flow-rate of 250 nL/min. LC was coupled with an Orbitrap Fusion mass spectrometer (Thermo Fisher Scientific, San Jose, CA, USA) with a spray voltage of 2.3 kV and capillary temperature of 275 °C. Full scan MS spectrum (m/z 300−1200) was acquired in the Orbitrap with a resolution of 60,000 (at 200 *m/z*) with an AGC target of 5x10e5. At Top Speed MS/MS option of 2 sec, the most intense ions above a threshold of 2000 counts were selected for fragmentation with higher-energy collisional dissociation (HCD) with normalized collision energy of 29, an AGC target of 1x10e4 and a maximum injection time of 200 msec. MS/MS data were collected in centroid mode in the ion trap mass analyzer (normal scan rate). Only charge states 2–4 were included. The dynamic exclusion was set at 30 sec. Where data-dependent acquisition^54^ was used to analyze the peptides, full scan MS (*m/z* 300−1100) was performed also in the Orbitrap with a higher resolution of 120,000 (at 200 m/z), AGC target set at the same 5x10e5. The difference is in the MS/MS though also performed in the ion trap, was with sequential isolation windows of 50 *m/z* with an AGC target of 3x10e4, a CID collision energy of 35 and a maximum injection time of 50 msec. MS/MS data were collected in centroid mode. For both acquisition methods, peak area was extracted from raw files by using our in-house software EpiProfile^55^. The relative abundance of a given PTM was calculated by dividing its intensity by the sum of all modified and unmodified peptides sharing the same sequence. For isobaric peptides, the relative ratio of two isobaric forms was estimated by averaging the ratio for each fragment ion with different mass between the two species.

**RNA isolation**. Total RNA extraction from epididymal sperm and EV pellets were done using the TRIzol reagent (Thermo Fisher) according to manufacturer’s protocol. For RNA extraction of PVN punches, the RNeasy Micro Kit was used according to manufacturer’s protocol (Qiagen).

**RiboTag mRNA immunoprecipitation**. To obtain actively translating mRNA, RiboTag mice were used as previously described^46^. Briefly, whole caput epididymal tissue were dounce homogenized in 1 ml supplemented homogenization buffer (50 mM Tris pH 7.5, 100 mM KCl, 12 mM MgCl_2_, 1% NP-40, 1mM DTT, 200U/mL RNasin (Promega), 1mg/mL heparin, 100 μg/mL cyclohexamide, protease inhibitor cocktail (Sigma)). Following centrifugation at 10,000g for 10 min, 800μl of the supernatant was incubated with 5μl of anti-HA.11 clone 16B12 antibody (Biolegend) for 4 h at 4°C. 400 μl of Dynabeads Protein G (Life Technologies) were washed with supplemented homogenization buffer and incubated with the supernatant-antibody complex overnight at 4°C. The next morning, bead-antibody-protein complexes were washed 3 times for 10 min with high salt buffer (50 mM Tris pH 7.5, 300 mM KCl, 12 mM MgCl_2_, 1% NP-40, 1 mM DTT, 100 μg/mL cyclohexamide). Immediately following washes, Qiagen Buffer RLT with beta-mercaptoethanol was added and the RNeasy protocol was followed according to manufacturer’s protocol to isolate RNA from the complexes.

**mRNA sequencing and analysis**. Total RNA from caput epididymal RiboTag and PVN punches were quantified on a NanoDrop 2000 spectrophotometer (Thermo Scientific). Libraries for RNA-seq were made using a TruSeq Stranded mRNA Sample Preparation Kit (Illumina) with 250ng RNA according to manufacturer’s protocol. All library sizes and concentrations were confirmed on a TapeStation 4200 (Agilent) and Qubit 3.0 Fluorometer (Thermo Fisher). Individually barcoded libraries were pooled and sequenced on an Illumina NextSeq 500 (75-bp single-end). Fastq files containing an average of 50 million reads were processed for pseudoalignment and abundance quantification using Kallisto (version 0.43.1)^56^. The transcriptome was aligned to the EnsemblDB Mus musculus package (version 79).

**Small RNA sequencing and analysis**. Small RNA libraries were constructed using the NEBNext Small RNA Library Prep Set for Illumina (NEB) on 200ng total RNA according to manufacturer’s protocol. All library sizes and concentrations were confirmed on a TapeStation 4200 (Agilent) and Qubit 3.0 Fluorometer (Thermo Fisher). Individually barcoded libraries were pooled and sequenced on an Illumina NextSeq 500 (75-bp single-end). Fastq files containing an average of 10 million reads per sample were aligned and quantified using miRDeep2 (version 2.0.0.8)^57^.

**Bioinformatics analyses:** All analyses were performed using R version 3.3.3 and Bioconductor version 3.4.

**Random Forests**. The R package randomForest^37^ was used to analyze histone mass spectrometry ratio data with the parameters ntree = 1000 and mtry = √*p* for classification analysis, based on calculation of *p* where *p* = total number of histone modifications identified. This approach ranks each histone modification by the percent decrease (MDA) to the model’s accuracy that occurs if the histone mark is removed, allowing for the identification of a histone code that discriminates between treatment groups. To estimate the minimal number of histone modifications required for prediction, ten-fold cross-validation using the ‘rfcv’ command was implemented through the randomForest package.

**Rank-rank hypergeometric overlap (RRHO)**. The R package RRHO was used to evaluate the degree and significance of overlap in threshold-free differential expression data between *in vivo* sperm and *in vitro* EV miR datasets^33^. For each comparison, one-sided enrichment tests were used on –log_10_(nominal p-values) with the default step size, and corrected Benjamini-Yekutieli p-values were calculated. Each pixel represents one miR comparison between the two datasets, with the degree of significance color-coded.

**Differential expression analysis**. The R package DESeq was used to perform pairwise differential expression analyses on RNA-seq datasets using the negative binomial distribution^58^. For PVN and RiboTag mRNA-seq, count data were filtered for at least 10 counts per gene across all groups, normalized, and dispersions were estimated per condition with a maximum sharing mode. Small RNA-seq data were filtered for greater than 2 counts in at least 3 samples across all groups, normalized, and dispersions were estimated per-condition using empirical values. Significance for all differential expression was set at an adjusted *P*-value<0.05. Heatmaps were generated using the R package gplots heatmap.2 function. All heatmaps are plotted as average Z scores per treatment group and arranged through hierarchical clustering of groups. Clusters of co-regulated differentially expressed genes were determined with the R package Stats using hierarchical clustering of genes (complete method) followed by ‘cutree’, k = 3.

**ClueGO**. Functional annotation analysis was performed on co-regulated differentially expressed gene clusters with the Cytoscape plug-in ClueGO^47^. ClueGO identifies enriched pathways using Gene Ontology (GO) terms and can reduce redundancy of GO terms that share similar genes by sorting into parent categories. For each cluster of differentially expressed genes, ClueGo was used to determine the enriched GO biological processes. Redundant terms were allowed to fuse with related terms that had similar associated genes. Networks of GO terms visualized using Cytoscape were linked using kappa statistics, with connecting nodes sized according to *P*-values corrected by Bonferroni step down. For each group of related GO terms, the leading group was determined by highest degree of significance. Top enriched groups of GO terms for each cluster were determined by the corrected group *P*-value.

**Statistics**. Corticosterone levels were analyzed by two-way ANOVA with time as a repeated measure. Corticosterone AUC, litter characteristics, testis weights, and gene expression data were analyzed by two-way ANOVAs. Outliers for HPA axis assessment were excluded at all time points and determined by data greater than two standard deviations away from the group mean or corticosterone levels greater than 150 ng/mL at the 120 min time point, indicating no stress recovery. Immunoblotting data were analyzed using one-way ANOVAs or two-tailed *t*-tests. Nanosight, and IVIS radiant efficiency were analyzed using two-tailed *t*-tests. Histone mass spectrometry ratio data were analyzed Mann-Whitney U tests. When appropriate, Bonferroni’s multiple comparisons or Student’s *t*-tests were used to explore main effects. Proportions of reproductive success were analyzed using chi-square tests. Significance was set at *P*<0.05.

## Acknowledgements

Research was supported by NIH grants MH108286, MH104184, MH099910, and ES028202 to T.L.B. and GM110174 to B.A.G. We thank D. Beiting for bioinformatics consultation; C. Howard, J. Fluharty, C. O’Donnell and H. O’Donnell for technical assistance; E. Pure’s lab for tissue culture assistance; and the Model Animal Research Center of Nanjing University for the Lcn5-Cre transgenic mice. Data for this study are available at the NCBI Gene Expression Omnibus (GEO) database, accession numbers GSEXXX.

## Author Contributions

J.C.C. and T.L.B. conceived the study and wrote the manuscript. J.C.C. performed most experiments with the help of B.M.N., K.E.M., and N.V.B. B.A.G. and N.V.B. assisted with histone mass spectrometry. J.C.C. analyzed the data with the help of B.M.N., E.J., and N.V.B.

